# Taxonomic notes on New Zealand orchids II: typification of *Nematoceras trilobus* Hook.f. and the names of four *Thelymitra* species described by William Colenso

**DOI:** 10.1101/2025.10.21.683774

**Authors:** Hayden Jones, Carlos A. Lehnebach

**Author notes:** Contribution notes: Both authors contributed equally to the research and writing of this article.

## Abstract

Names for five species of Orchidaceae from New Zealand described by Joseph Dalton Hooker and William Colenso are here typified. This includes lectotypes for *Nematoceras trilobus* Hook.f. (the basionym of *Corybas trilobus* (Hook.f.) Rchb.f.), *Thelymitra nemoralis* Colenso*, Thelymitra nervosa* Colenso and *Thelymitra purpureo-fusca* Colenso. We also report finding the holotype of *Thelymitra concinna* Colenso at WELT and discuss synonymisation of *T. decora* Cheeseman with *T. nervosa*.

## Introduction

The taxonomic description of the New Zealand (NZ) Orchid Flora began more than 250 years ago when Joseph Banks and Daniel Solander landed in NZ in 1769 and collected the first orchid specimens. Since then, several visiting and resident naturalists explored NZ, collecting biological specimens that were mostly sent to overseas institutions such as the Royal Botanic Gardens at Kew. It was there where the botanist Joseph Dalton Hooker (1817–1911) described many of NZ’s most common orchids (∼ 20 species, St. George 2008a; IPNI 2025). Towards the late 1880s, however, there was a shift in this trend when Rev. William Colenso (1811– 1899), a Christian missionary and naturalist based in the Hawke’s Bay (North Island), began to write his own species descriptions. Overall, Colenso described 36 orchid species (St. George 2008a; IPNI 2025). These species were mostly published between 1880 and 1890 in the *Transactions and Proceedings of the New Zealand Institute*. After publication, Colenso would ship representative material, likely duplicates, of his new orchid species to Kew (St. George et al. 2009). However, this was not always the case (see Lehnebach & Jones 2024) and Moore (1976), who prepared the most recent family-level revision of NZ orchids, was unable to locate Colenso’s original material at Kew for several orchid species.

Tracing the whereabouts and identity of Colenso’s type specimens can be a complex task. Colenso claimed that “*he did not keep an herbarium*” and believed that “*keeping specimens for species described from cultivated material was unnecessary*” (Colenso 1886; Hamlin 1971; Large et al. 2020). Further confusion with the status and origin of his specimens was created by Thomas F. Cheeseman (1845–1923), who explicitly challenged many of Colenso’s species and relabelled Colenso’s herbarium before it reached WELT. This was followed by a re-organisation, mostly mounting, of Colenso’s material once at WELT. For further details on Colenso’s herbarium see Lehnebach and Jones (2024) and references therein.

Currently, it is accepted that more than 115 indigenous species of orchids occur in NZ (Allan Herbarium 2024); this number is likely to increase as the taxonomic status of numerous entities known by tag-names only (see de Lange et al. 2024) is being assessed using genetic and morphological data by the authors of this article and collaborators. Many of these “potentially new species” are part of highly variable species or species aggregates of *Corybas*, *Pterostylis* and *Thelymitra* (de Lange et al. 2024), NZ’s largest orchid genera.

In preparation for the publication of taxonomic novelties in NZ *Corybas* and *Thelymitra* resulting from these studies, solving the taxonomic and nomenclatural status of the names of five orchid species described by J. D. Hooker and W. Colenso is necessary. To this end, in this article we designate lectotypes for *Nematoceras trilobus* Hooker.f. (i.e. the basionym of *Corybas trilobus* (Hook.f.) Rchb.f.), *Thelymitra concinna* Colenso, *T. nemoralis* Colenso, *T. nervosa* Colenso, and *T. purpureo-fusca* Colenso.

## Methods

We searched for Hooker’s and Colenso’s original material at herbaria in NZ (i.e., AK, CHR, WELT) and overseas (i.e., K, L, MEL, NSW, P, S). We also investigated Colenso’s archival collection at the Museum Theatre Gallery Hawkes Bay (MTG) (Napier, New Zealand). Material lodged in overseas herbaria was loaned or accessed via online portals such as the Atlas of Living Australia (2023), GBIF.org (2023), JSTOR Global Plants database (JSTOR 2025) or correspondence with curators and collection managers from these institutions. Herbarium acronyms follow Thiers (2025), and typification follows the guidelines outlined in the International Code of Nomenclature (ICN) (Turland et al., 2018).

Both, Dr Brian Molloy and Dr Mark Clements have studied and confirmed the identity of some of the material we studied and annotated them as type material. Their typification, however, was not effective as it was never published. When the specimens they selected as types aligned with our conclusions, we designate them as lectotypes.

## Results

### Nematoceras

*Nematoceras trilobum* Hook.f., Bot. Antarct. Voy. II. (Fl. Nov.-Zel.) Part I, 250 (1853) – as *triloba*.

**Type collection**: East Coast and interior [North Island, New Zealand], [William] *Colenso*.

**Lectotype**: (designated here) K000364466! (Figure 1, arrow)

**Figure 1.**
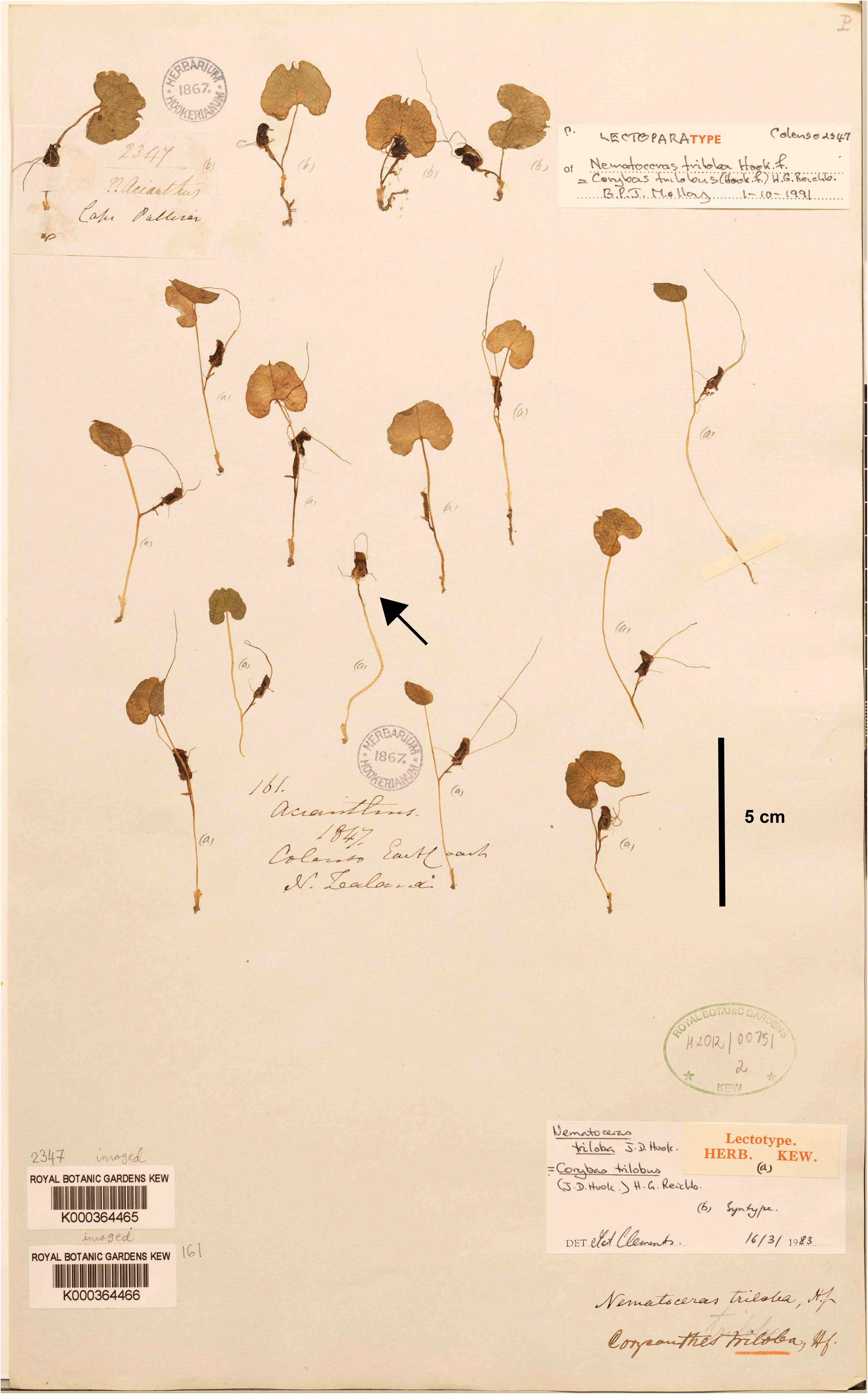
Lectotype of *Nematoceras trilobum* Hook.f. (arrow) collected by William Colenso at Kew Herbarium (K000364466). © RBG Kew.

**Notes**: The original material used by Joseph Dalton Hooker (1817–1911) for the description of *Nematoceras trilobum*, the basionym of *Corybas trilobus* (Hook.f.) Rchb.f., was collected by William Colenso between 1841 and 1850 and sent to Sir William Jackson Hooker (1785–1865) at Kew. This material is now part of the Herbarium Hookerianum and corresponds to three gatherings which can be identified by Colenso’s numbers associated with them, 16 (K000364467!), 161 (K000364466!) and 2347 (K000364465!). The first gathering consists of fruiting specimens and the last two of flowering specimens, which are particularly useful because in *Corybas* identification to species level is practically impossible if no flowers are present. The last two gatherings are mounted on the same sheet and annotated “*Nematoceras triloba* Hook.f.” and “*Corysanthes triloba* Hook.f.” (a later synonym of *N. trilobum*) (Figure 1).

At the top of this sheet there are four specimens associated with a hand-written label, not in Colenso’s hand, that reads “2347 ??*Acianthus* - Cape Palliser”. This note refers to the number Colenso assigned to this collection when preparing a list of the specimens to be sent to Kew, Colenso’s tentative identification, and the locality where the plants were collected. All four specimens are annotated *“(b)”* and a modern label, signed by B.P.J. Molloy in 1-10-1991, identifies this gathering as “lectoparatype”. The second gathering, in the centre-lower part of the sheet, consists of 12 individuals. A note, not in Colenso’s hand, reads “161. *Acianthus*. 1847. Colenso East Coast. N[ew]. Zealand”. As above, this information refers to the number Colenso assigned to the gathering, his tentative identification, the year when the material arrived at Kew (St. George et al. 2009) and collection locality. The latter detail, however, does not fully match Colenso’s notes. In his list, Colenso stated that the collection 161 was found “*in flower, in shaded damp spots, wood, with Nos. 154 & 159*”. In the same list, the collection 154 is said to be “*from forest, hills, between Wareama* [Whareama, about 40K east from Masterton, in the Wairarapa, North Island], *& the head of Wairarapa Valley*”. Collection 159 was found in “*woods, with 154*”. Thus, the correct origin of 161 is between Whareama & the head of Wairarapa Valley. All 12 specimens are annotated “*(a)*”. A modern label, signed by Mark Clements in 16-03-1983, assigns these specimens to lectotype and those labelled *(b)* to syntypes. To the best of our knowledge, Clements’ typification has never been effectively published.

Closer examination of both gatherings shows the flowers of each collection are morphologically distinct from each other (Figure 2, A and C), confirming they belong to two different taxa. The most obvious differences are the colour of the labellum and the distal end of the dorsal sepal, length of the lateral petals and the lower margin of the labellum. This means Hooker’s description was based on a mixture of taxa and that lectotypification is necessary following Article 9.11, Article 9.12 of the ICN (Turland et al. 2018). From these two gatherings, the specimens annotated “*(a)*” and already selected by Clements as lectotype correspond most closely with the original description of *Nematoceras trilobum*, specifically in features such as leaf width, and length of the perianth, lateral sepals and petals. Clements’s selection, therefore, follows well Article 9.14 of the ICN (Turland et al. 2018). Furthermore, this material aligns with the current use of this name (see Rolfe 2016 and Figure 1 therein), which will prevent further confusion around the application of this name and species boundaries within this already difficult species complex. The material annotated *(b)* and labelled by Molloy as “*lectoparatype*” it is likely to correspond to the taxonomically undetermined entity currently known as *Corybas* aff. *trilobus* (CHR 534742; Trotters Gorge), which is also found in the Wairarapa and east coast of the lower North Island.

**Figure 2.**
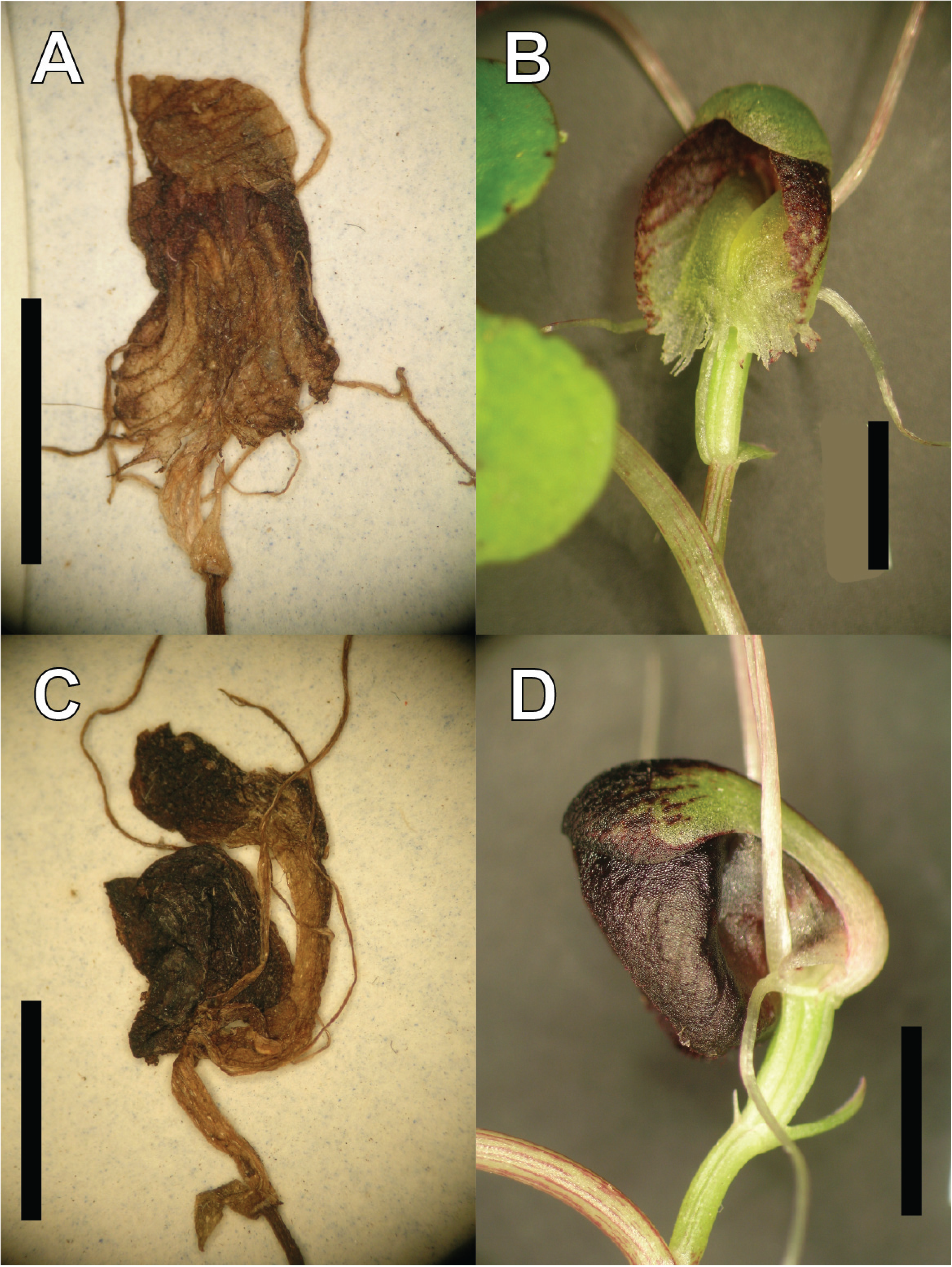
Close-up view of herborised and fresh flowers of *Corybas trilobus* (Hook.f.) Rchb.f. (syn*. Nematoceras trilobum* Colenso) (A, B) and possibly *Corybas* aff. *trilobus* “Trotters Gorge” (C, D). Material from K000364466 (A), WELT SP104184 (B), K000364465 (C), WELT SP110643 (D). Scale bar = 5mm. A, C: © RBG Kew. B, D: © Te Papa.

Lastly, Cheeseman (1906, 1925) and Moore (1976) incorrectly considered *C. hypogaeus* (Colenso) Lehnebach synonymous with *C. trilobus*. Cheeseman did not provide support for his decision, but Moore (1976) did for hers, stating that Colenso’s (1884) description “*applies mostly entirely to C. trilobus*”. She further suggested the description of *C. hypogaeus* was based on a mixed collection that also included material of *Corybas cryptanthus* Hatch, a mycoheterotrophic species that, as *C. hypogaeus*, normally grows half-buried in the leaf litter. She based this idea on Colenso’s chosen epithet, i.e. “*hypogaea*”. Colenso’s original material of *C. hypogaeus* is at Kew (K000942561!, K000942562!), and its study confirms that both Cheeseman and Moore were mistaken as it does not include material of *C. cryptanthus* or *C. trilobus*.

### Thelymitra

#### *1. Thelymitra concinna Colenso, Trans. & Proc. New Zealand Inst*. 20: 207 (1887 [1888])

**Type collection**: Open country near the east bank of the River Mohaka, north of Napier, 1884, Mr. *A. Hamilton*

**Holotype**: WELT SP024275! (Figure 3)

**Figure 3.**
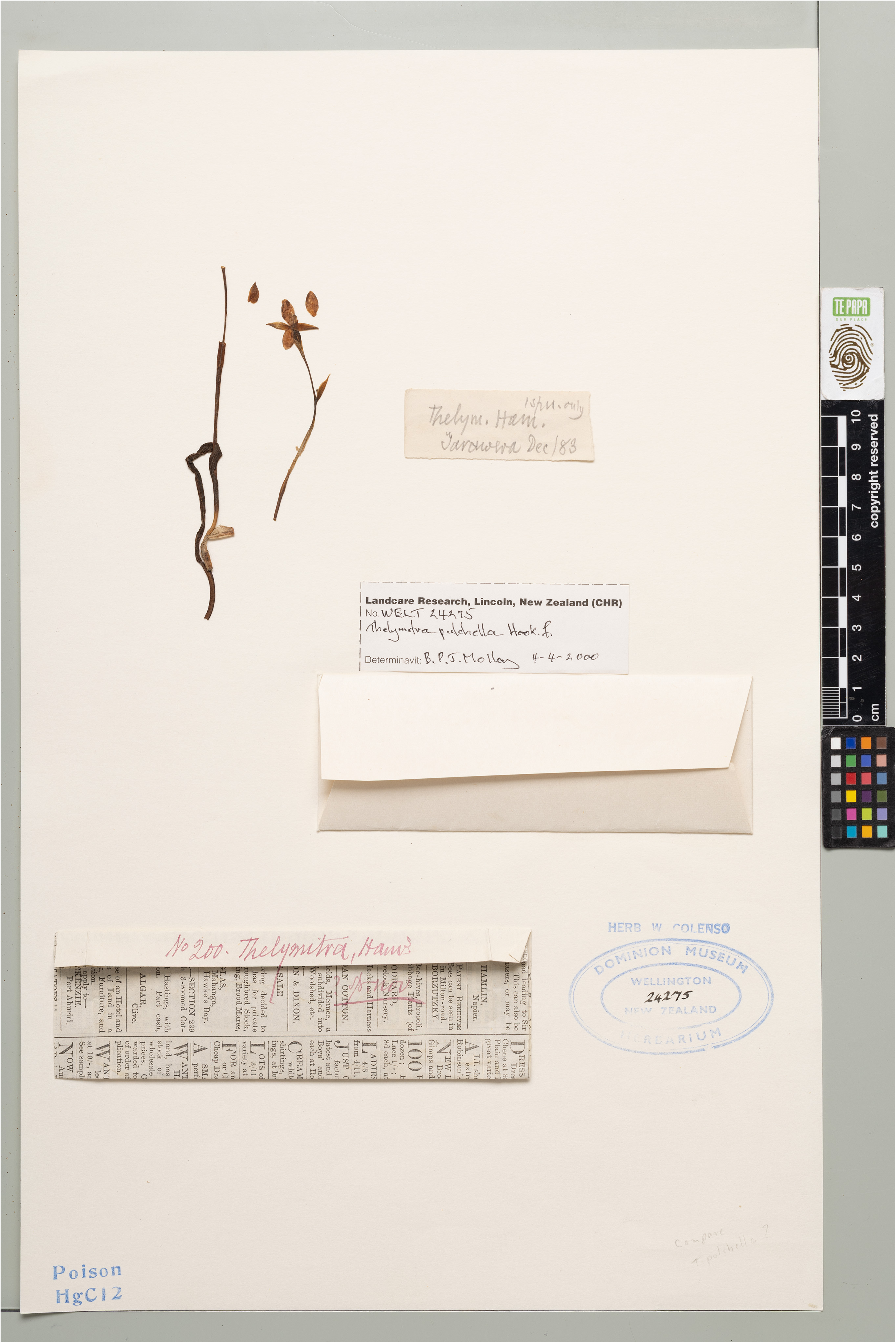
Holotype specimen of *Thelymitra concinna* Colenso at WELT Herbarium (SP024275). © Te Papa.

**Notes**: In the protologue Colenso mentioned he used a single specimen for the description of this species given to him by Mr. A. [Augustus] Hamilton (1854–1913), who collected it from the east banks of the Mohaka River (Hawke’s Bay, North Island). Colenso deferred publication of the species for a few years, hoping Hamilton would obtain more specimens, which did not happen.

Currently, *T. concinna* is considered a synonym of *T. pulchella* Hook.f. (Allan Herbarium 2024), a species described in 1853, but the reason for this decision has never been provided. Cheeseman (1906) did not include *T. concinna* in his *Manual of the New Zealand Flora,* only stating this species was unknown to him. The name did not feature at all in the second edition of the *Manual* (Cheeseman 1925). Moore (1976) was the first one to synonymise *T. concinna* with *T. pulchella,* but again no support was given for this decision. Moore only stated she was unable to find the type of *T. concinna*.

While studying the *Thelymitra* collection at WELT we found a *Thelymitra* specimen inside a newspaper envelope that we believe is Hamilton’s collection (Figure 3). The envelope is attached to the herbarium sheet WELT SP024275!. Writing on it, in Colenso’s hand, reads “*No. 200. Thelymitra Ham.*[Hamilton]*;sp nov*”. A second note inside the envelope, also in Colenso’s hand, reads “*1 spn. only; Thelym. Ham.*[Hamilton]; *Tarawera Dec/83*”. Tarawera (Hawke’s Bay, North Island) is a settlement along the Mohaka River, and within the type locality indicated in Colenso (1888). This evidence confirms the specimen WELT SP024275 is the holotype of *T. concinna* Colenso.

Following Moore’s opinion, NZ orchidologist Brian Molloy re-identified this material as *T. pulchella* and attached an identification slip to the sheet on 4/4/2000. However, upon closer examination of the flower we believe this material does not belong to *T. pulchella* as it lacks the flattened and fimbriate column arms, and the reduced, in-turned post anther lobe that are diagnostic for *T. pulchella* (see Figure 4 A, B). Colenso’s description and Hamilton’s collection resembles most closely *T. hatchii* L.B.Moore, differing only in the colour of the cilia (i.e., reddish), or, less likely, *T. formosa* Colenso. However, Colenso should have been familiar with this latter species since he collected it in 1882 and described it in 1883.

**Figure 4.**
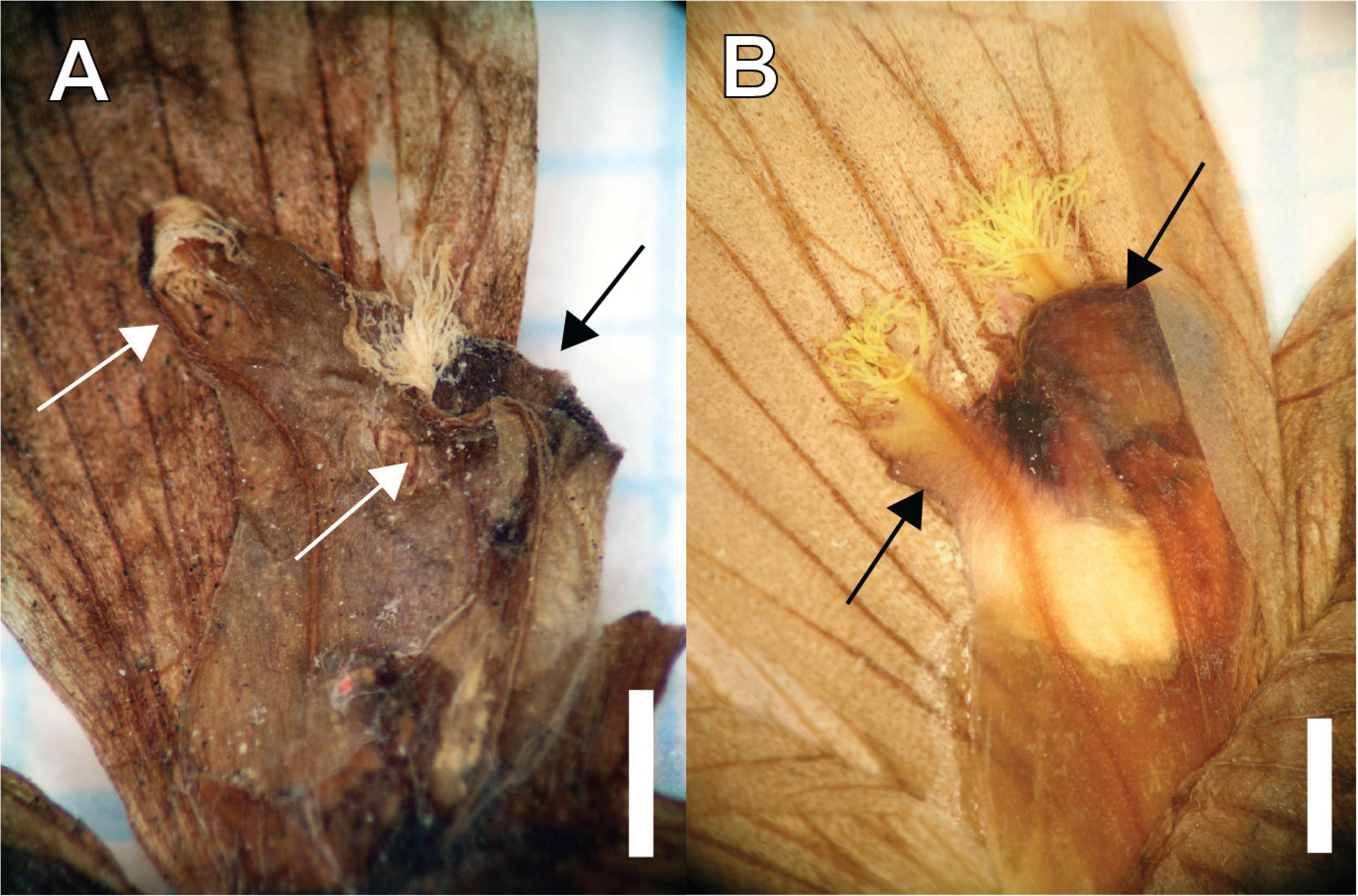
A detailed view of the column of *Thelymitra concinna* Colenso (A) and *T. pulchella* Hook.f. (B) showing diagnostic column features (arrows). Material from WELT SP024275 and WELT SP089994, respectively. © Te Papa.

*Thelymitra hatchii* is a highly variable species due in part to being an allopolyploid hybrid between *T. formosa* and *T. longifolia* (Molloy & Dawson 1998, Jones et al., 2025). This variability includes an uncommon pink-ciliated form (Figure 5) which we believe is what Colenso described as *T. concinna*. Following Articles 11.1 and 11.4 of the ICN (Turland et al., 2018), *T. hatchii* may have been nomenclaturally superfluous at the time of publication, with *T. concinna* having priority given its earlier publication date and legitimacy now that the holotype has been found. Given the obscurity of the name *T. concinna* however, and *T. hatchii* having widespread use and recognition, it may be a good case for name conservation following Articles 14.1 and 14.2 of the ICN (Turland et al., 2018). An application to the ICN for conservation of the name *T. hatchii* is currently in preparation by the first author.

**Figure 5.**
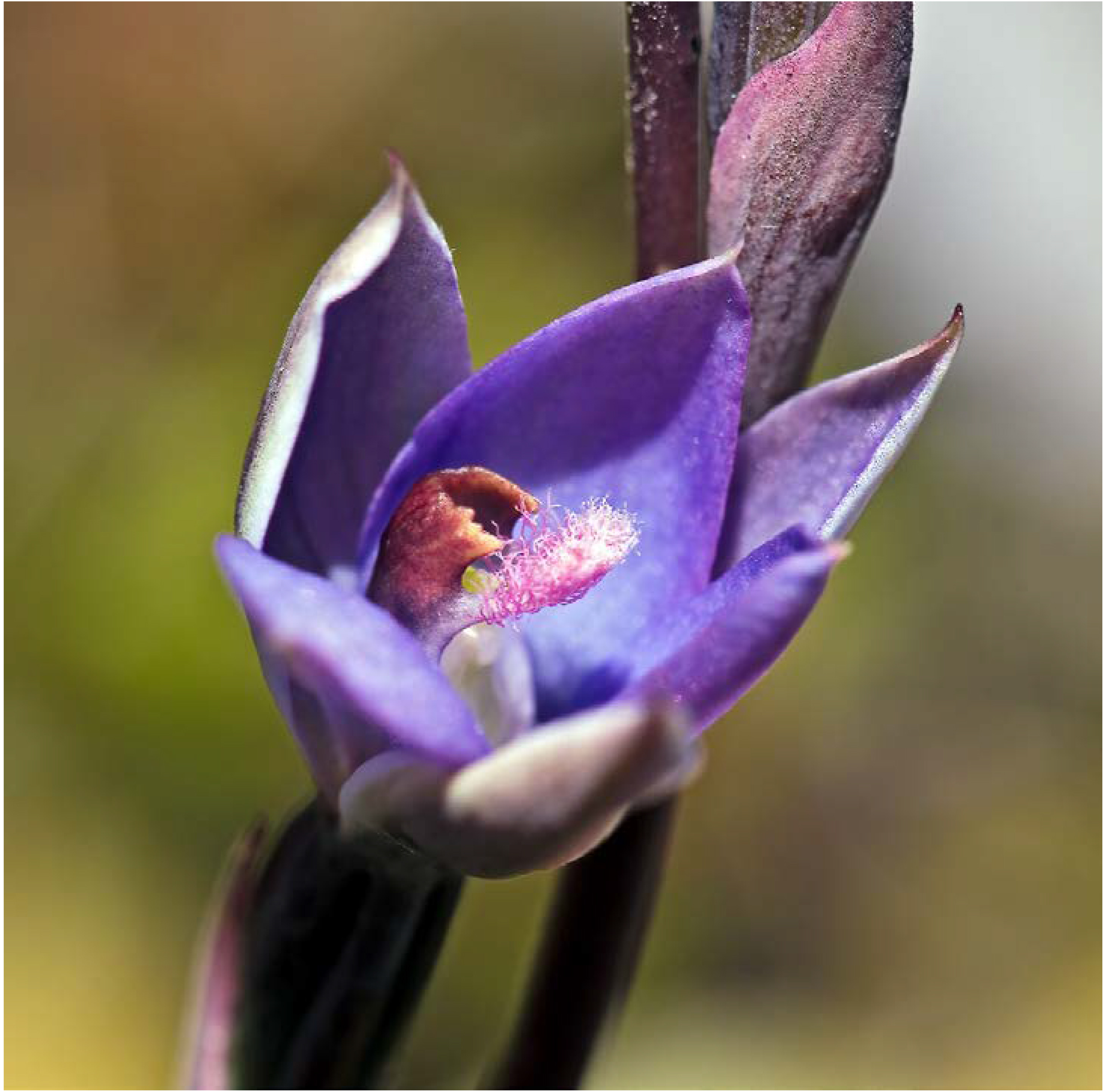
The flower of a pink ciliated *Thelymitra hatchii* L.B.Moore, which we believe is the phenotype Colenso described as *T. concinna* Colenso. Photo Credit: Jeremy Rolfe.

#### *2. Thelymitra nemoralis Colenso, Trans. & Proc. New Zealand Inst*. 17: 249 (1884 [1885])

**Type collection**: Dry *Fagus* [*Nothofagus*] Seventy-mile Bush, County of Waipawa, *W*.*C*. [William Colenso], 1881–83.

**Lectotype**: (designated here) K000827543! (Figure 6, specimen A)

**Figure 6.**
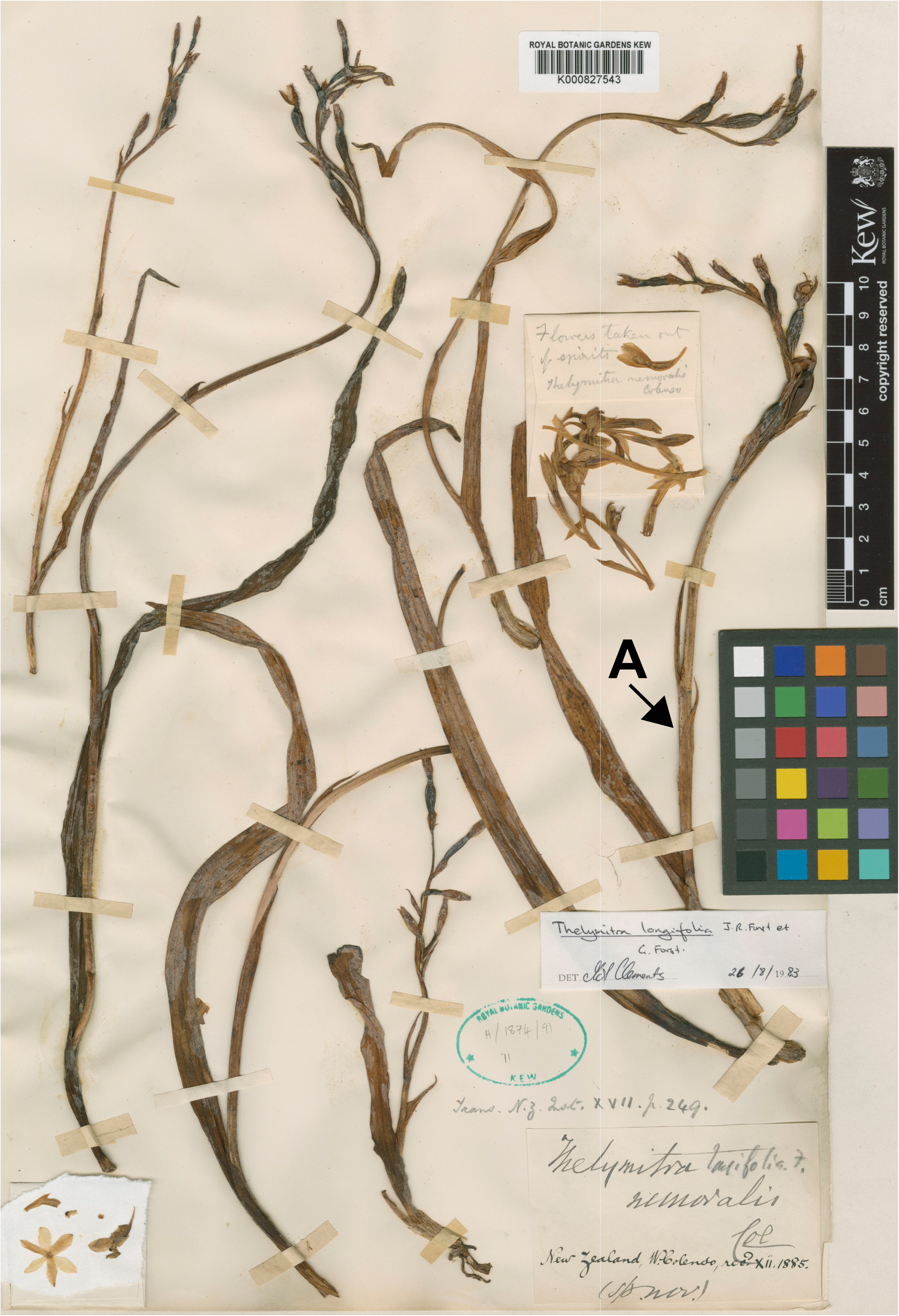
Lectotype specimen of *Thelymitra nemoralis* Colenso (specimen A) at Kew Herbarium (K000827543). Image copyright © RBG Kew

**Notes**: Colenso sent samples of *T. nemoralis* to Kew as both dried and spirit collections. Evidence for this appears in correspondence with J. D. Hooker dated 14^th^ October 1885 (St. George et al., 2009). At Kew, all Colenso’s material was laid together on sheet K000827543! which includes his original handwritten label with the species name [*Thelymitra nemoralis* Col (*sp*.*nov*.)]. A pencil annotation, not in Colenso’s hand, adds to Colenso’s label “*longifolia*. *f*.”. Two different notes, again not in Colenso’s hand, indicate when the material was received at Kew (i.e. XII – 1885) and where the species description was published (i.e. *Trans. N.Z. Inst*: XVII. p.249). A small envelop attached to this sheet contains flowers and is labelled in pencil “*Flowers taken out of spirits Thelymitra nemoralis Colenso*”. All this evidence confirms this is Colenso’s original material and a perfect candidate for typification. We select specimen A as the lectotype for *T. nemoralis*.

A determinative slip added by Mark Clements in 26/08/1983 identifies the material on K000827543 as *T. longifolia*. Synonymisation of *T. nemoralis* under *T. longifolia* has been widely accepted in NZ since the early 1900s (e.g. Cheeseman 1906) and has remained so since (see Allan Herbarium 2024). However, no reason has ever been provided for this decision. The genetic and morphological diversity within *T. longifolia* is currently under study by the first author and changes to this position are unjustifiable before that work is completed.

#### *3. Thelymitra nervosa Colenso, Trans. & Proc. New Zealand Inst*. 20: 207 (1887 [1888])

**Type collection**: High lands base of Mount Ruapehu (Tongariro Range), County of East Taupo, [unknown collector], 1879.

**Lectotype**: (designated here) K000827538! (Figure 7)

**Figure 7.**
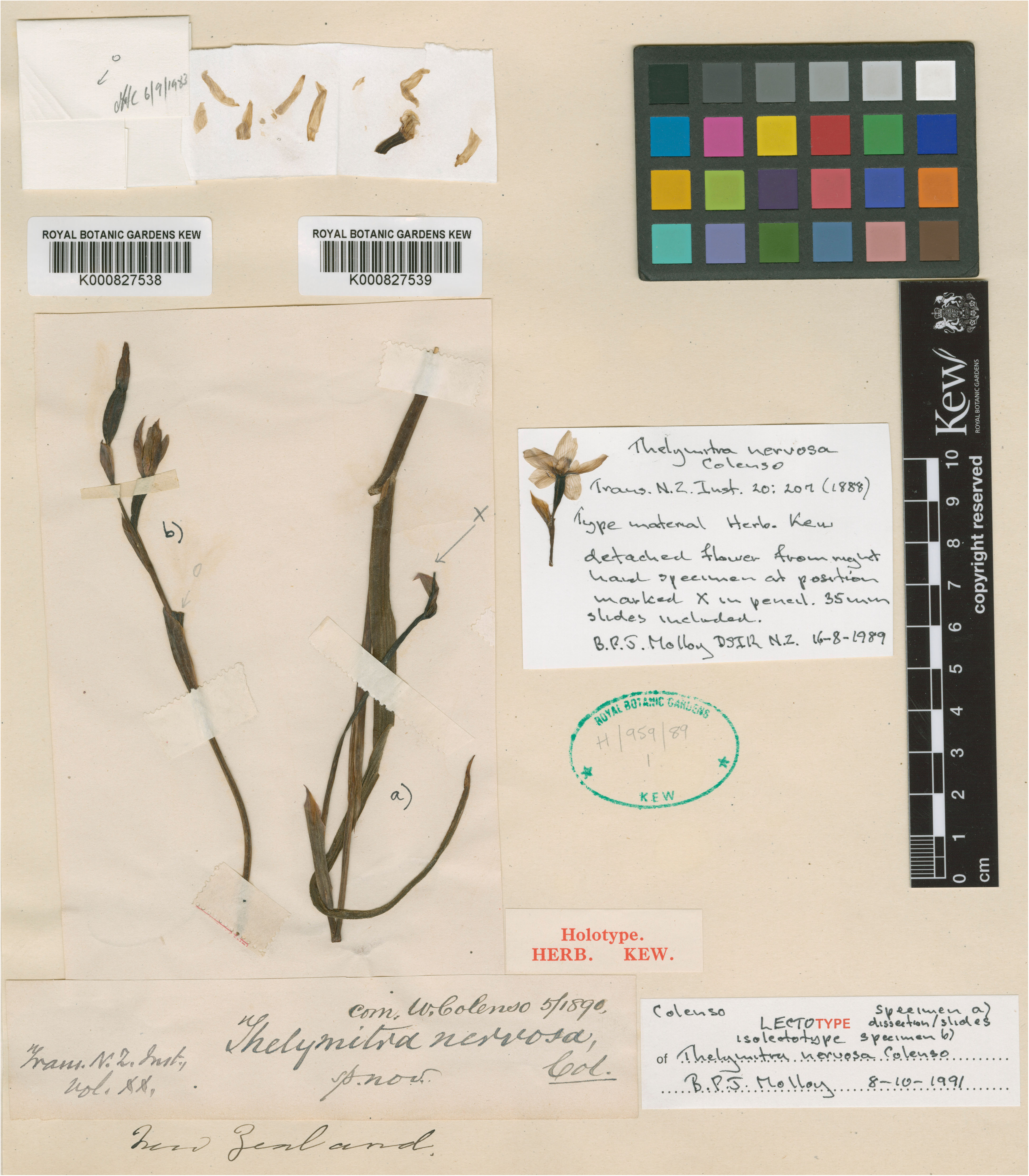
Lectotype specimen of *Thelymitra nervosa* Colenso at Kew Herbarium (K000827538). © RBG Kew

**Notes**: The only material that can be associated with this name is found at Kew and consists of two specimens (K000827538! and K000827539!), both attached to the same herbarium sheet. A label, in Colenso’s hand, identifies the material as “*Thelymitra nervosa*, Col. *sp. nov*” and indicates where it was published (i.e. *Trans. N.Z. Inst*; vol. XX). Two other annotations, not in Colenso’s hand, indicate country of origin (i.e. New Zealand) and date when the material reached Kew (i.e. May 1890). Unlike with many other specimens, Colenso made no mention of *T. nervosa* in his letters to Hooker or included it in the lists he normally prepared for each specimen shipment. Given that this material reached Kew in May 1890, it is likely Colenso sent these two samples with a shipment mentioned in correspondence to Sir William Turner Thiselton-Dyer (1843–1829, assistant director at the Royal Botanic Garden) dated 4^th^ March 1890 (see St. George et al., 2009). In this letter Colenso said: “… *I have concluded my set task (begun in June/88!) of putting up spns.* [specimens] *for Kew. These will go by the first steamer hence … of phaenograms and Ferns, 70* [specimens]*, or more – nearly all of these last spns.* [specimens] *are sps. novae, and described by me in Trans. N.Z. Inst.*”.

There are two modern labels fixed to this herbarium sheet, both by B.P.J. Molloy. The first one, dated 16-08-1989, indicates the specimen belongs to *T. nervosa* and that is “*type material*”. The second label is dated 8-10-1991, and indicates specimen a), the floral dissection (shown in Figure 8 A) and slides represent his chosen lectotype (K000827539!), and the specimen b) (K000827538!), on the left-hand side, his chosen isolectotype. We agree with Molloy’s selection and formally designate here the specimen K000827539! as the lectotype of *T. nervosa*.

**Figure 8.**
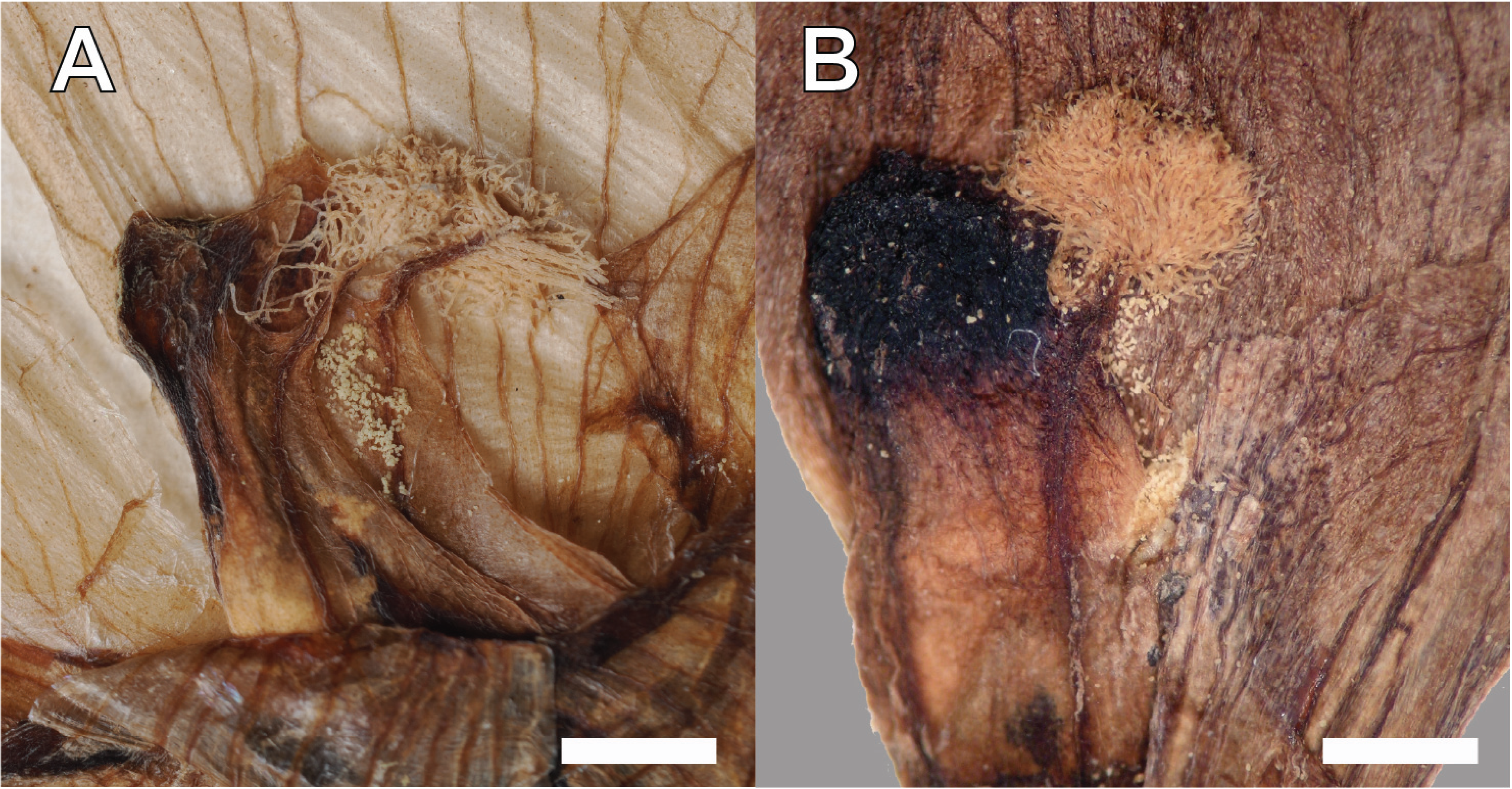
Columns of *Thelymitra nervosa* Colenso (A) and *T. decora* Cheeseman (B). Material from K000827538 and WELT SP18575, respectively. Photos by A. Schuiteman, Royal Botanic Gardens, Kew (A) and C. Lehnebach, Te Papa. Scale bar = 1 mm.

Application of the name *T. nervosa* and its status have been regularly debated among the botanical community and until very recently the name had been excluded from many taxonomic works. For instance, Cheeseman (1906) did not include *T. nervosa* in his *Manual* and stated that *T. nervosa*, along with *T. concinna*, were unknown to him. No reference to *T. nervosa* is found in Cheeseman (1925). Almost 50 years later, Moore (1976) considered *T. nervosa* as a species of uncertain placement and stated …“*Colenso described a number of spp. in the years 1884-1890 and several of these names are not supported by any known specimens. Most* [of these] *can be placed with varying degrees of confidence in the synonymy of accepted spp. but one remains unresolved; this is T. nervosa*…”

At some point, within the last 30 years, the name *T. nervosa* became widely used in NZ and *T. decora* Cheeseman, a species described in 1906, has been synonymised under it. The first published record of this change that we were able to find is a checklist by St. George (1997). This checklist included the name *T. nervosa* followed by the note “*it has been named T. decora”*. A disclaimer at the beginning of the checklist, however, states the list “*should not be regarded as a definitive list of the New Zealand species*” and that it represents “*a subjective and composite view, gleaned by the editor* [i.e. St. George] *from reading, conversation and observation*”. Since then, synonymisation of *T. decora* with *T. nervosa* has been widely accepted nationally (e.g. St George et al 2001, Dawson et al 2007, Breitwieser et al 2012) and internationally (e.g. POWO 2025). Despite our efforts, we have not found published justification for this decision.

Assuming these two names are synonymous, *T. nervosa* has priority over *T. decora* given its earlier publication date. However, one major issue with this position, is the lack of characters in the description of *T. nervosa* matching the material circumscribed as *T. decora* by Cheeseman (1906). In fact, particular characters that are diagnostic for *T. decora* (i.e. lateral petals spotted, column margin denticulate, back of column minutely warted) are not mentioned in the protologue of *T. nervosa* – which instead described the petals and sepals as being “*much veined*” (the presence of conspicuous veins on the perianth is the reason for Colenso’s chosen epithet *nervosa*) and the column “*largely bifid, each lobe 1-notched and incurved*”. Differences between the columns of both species can be appreciated in Figure 8 A & B. Both are from original material studied by Colenso and Cheeseman, respectively. Furthermore, none of the diagnostic features of *T. decora* are visible from an illustration at Kew (K001568512, Supplementary Figure 1) believed based on Colenso’s original material (i.e. K000827538!) (Dr André Schuiteman - Kew Herbarium, *pers. comm*.) and identified as *T. nervosa* by M.A. Clements. In view of these findings, our recommendations are to exclude this name from the synonymy of *T. nervosa* and re-establish the use of *T. decora*.

This action, however, will cast doubt on the application of the name *T. nervosa*. There are three taxa of *Thelymitra* in NZ with obviously “*veined*” flowers: *T. cyanea* Lindl. *ex* Benth., *T*. × *dentata* L.B.Moore and *T. pulchella* Hook.f.. Of these three, both *T*. × *dentata* and *T. pulchella* have column arms with cilia (which are mentioned in Colenso’s description of *T. nervosa*) but only the cilia of *T*. × *dentata* could be described as “*largely plumose*”. Cilia are always sparse in *T. pulchella*. Moore (1968) also noticed the similarity of *T. nervosa* with *T*. × *dentata* when describing the latter, but she was unable to find Colenso’s original material and compare them. *Thelymitra* × *dentata* is a sterile hybrid between *T. longifolia* and *T. pulchella* (Dawson et al. 2007) which is uncommon but has been reported from *T. nervosa*’s type locality (see https://www.inaturalist.org/photos/457758252). A thorough morphological assessment of fresh and historical collections will be necessary to uncover the existence of plants matching Colenso’s description and original material of *T. nervosa*. This should also include surveys to the type locality near Mount Ruapehu and genetic studies to detect potential hybridisation (see Jones et al. 2025).

#### *4. Thelymitra purpureofusca Colenso, Trans. & Proc. New Zealand Inst*. 17: 249 (1884 [1885])

**Type collection**: In *Fagus* [*Nothofagus*] with the preceding species [*T. nemoralis*], but usually higher up, 1881–83, *W*.*C*. [William Colenso]

**Lectotype**: (designated here) K000827535! (Figure 9, specimen A)

**Figure 9.**
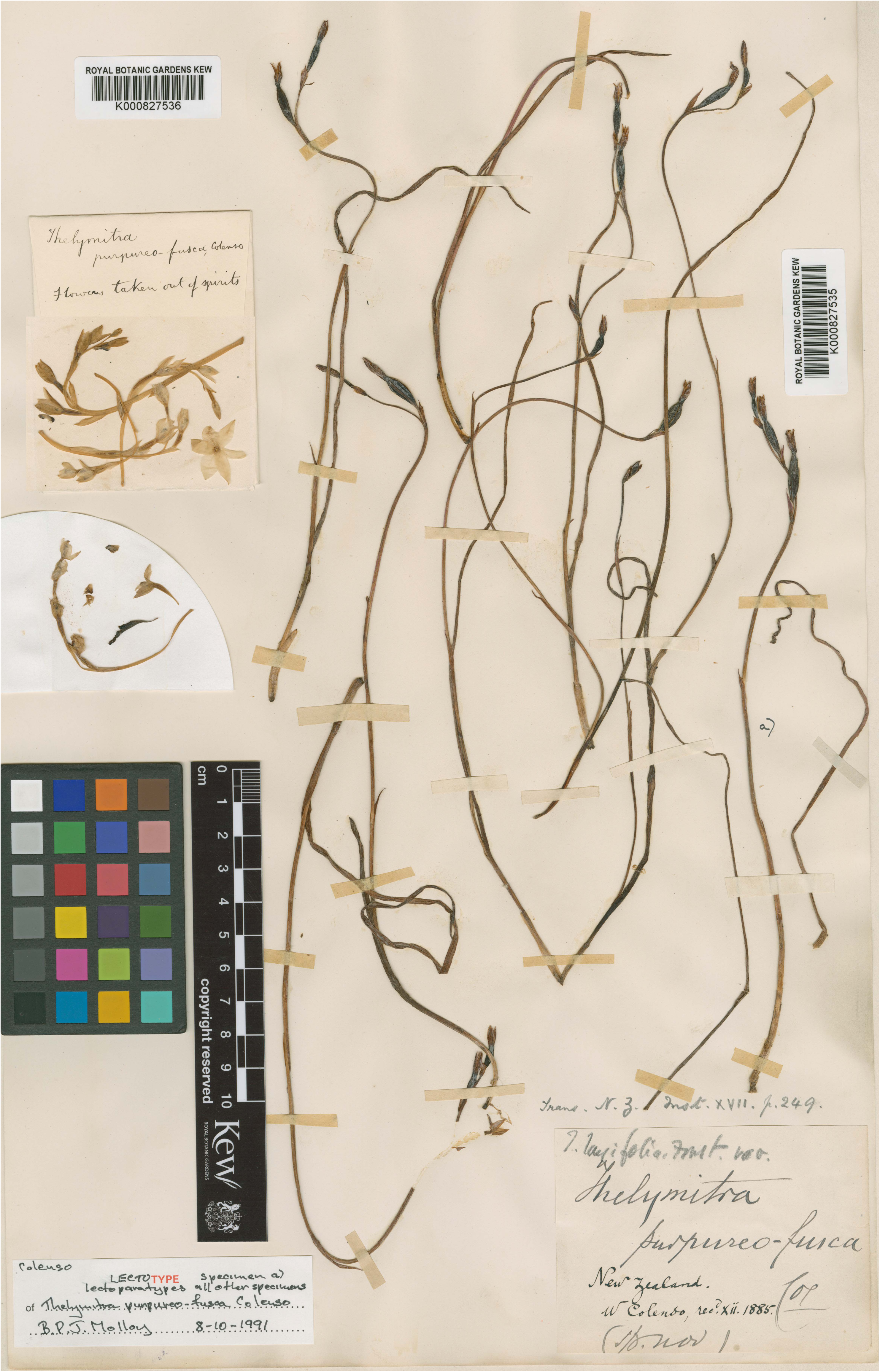
Lectotype of *Thelymitra purpureofusca* Colenso (specimen a) at Kew Herbarium (K000827535). © RBG Kew.

**Notes**: As he did with *T. nemoralis*, Colenso sent dried and preserved material of his new species to Kew, and this was recorded in the correspondence with J. D. Hooker on 14 October 1885 (St. George et al. 2009). At Kew the material has been affixed to a single herbarium sheet; the dry-pressed material has the accession number K000827535! and that taken out of spirits K000827536! A label, in Colenso’s hand, reads “*Thelymitra purpureo-fusca* Col (*sp*. *nov*)”. Two other annotations, one in ink and the other one in pencil, indicate the origin and collector (*New Zealand*. *W Colenso*), date when it was received (xii. 1885), and the species publication details (*Trans. N.Z. Inst*. XVII. p.249). None of these are in Colenso’s handwriting. Based on the information provided above we believe the material on this sheet is the best candidate for typification. A similar conclusion was reached by Brian Molloy who, in 8-10-1991, selected “*specimen a*” on K000827535! as lectotype but, to the best of our knowledge, his typification was never published.

Historical treatments (Cheeseman 1906, 1925; Hatch 1952, Moore 1976) have placed *T. purpureofusca* in synonymy with *T. longifolia*. A pencil annotation on the herbarium sheet K000827535! that reads “*T. longifolia* Forst. var.” seems to be in line with these authors. Most recently, however, the taxonomic aggregator POWO (2025) lists *T. purpureofusca* as an “accepted species” occurring in the North Island of NZ. A species list compiled by St. George (2008b) is used to support that status. Curiously, this name does not appear in the latest version of the Checklist of the NZ flora - seed plants by the Allan Herbarium (2024).

Genetic and morphological studies by the first author are looking into many of Colenso’s species nowadays synonymised with *T. longifolia*, including *T. purpureofusca*. Until those studies are completed, we recommend including *T. purpureofusca* in the Checklist of the NZ flora prepared by the Allan Herbarium (CHR) but as a synonym of *T. longifolia*.

## Acknowledgements

We would like to thank the curators, collection managers and collection technicians from AK, CHR, E, K, MTG and WELT from their help accessing non-digitised collections. We also appreciate comments and suggestions by two anonymous reviewers and Dr Barry Sneddon and Dr Lara Shepard on an earlier draft of this manuscript.

