## Supplementary Figure 1 for "Taxonomic notes on New Zealand orchids II: typification of *Nematoceras trilobus* Hook.f. and the names of four *Thelymitra* species described by William Colenso"

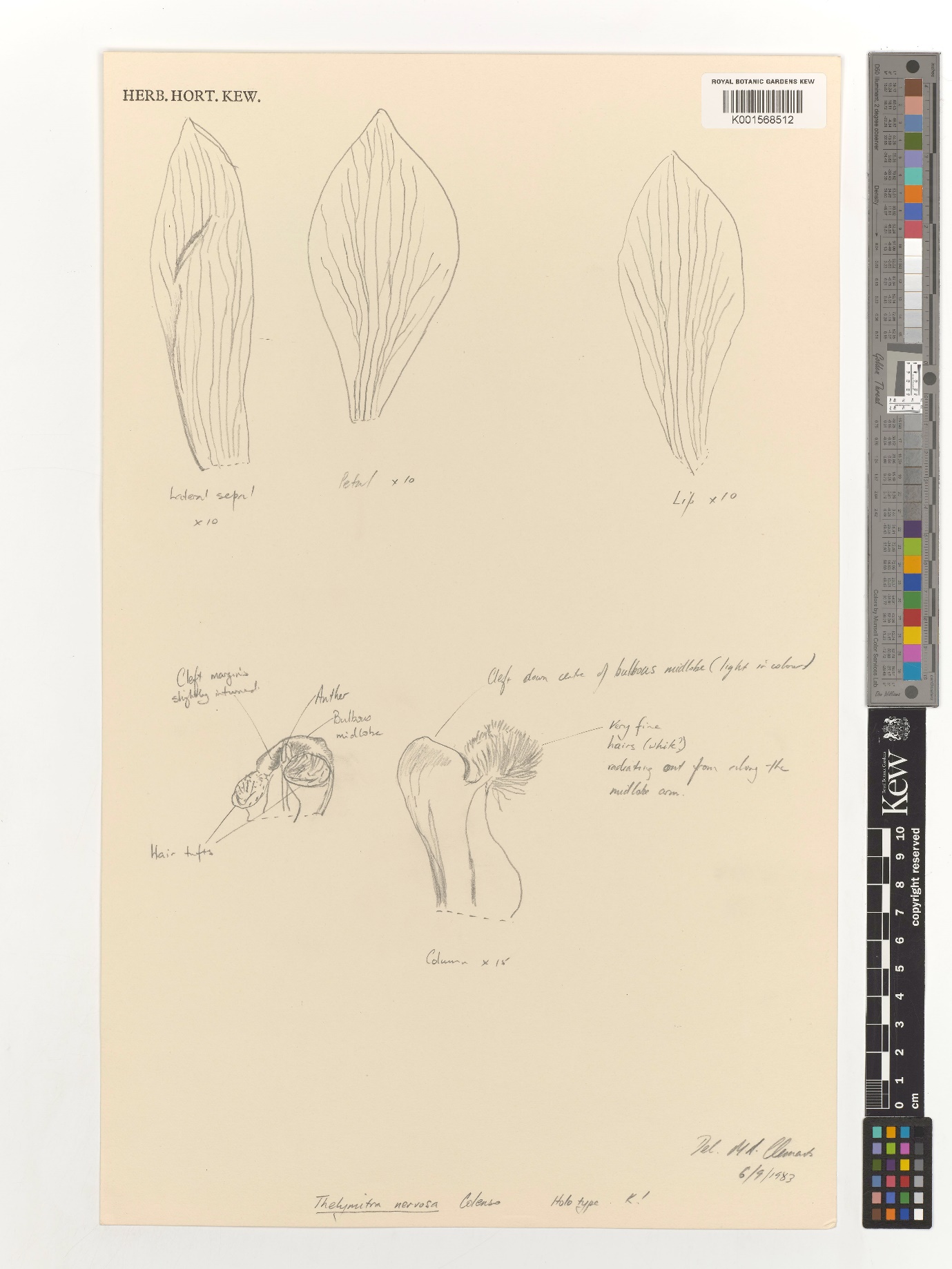

Supplementary Figure 1: *Thelymitra venosa* Colenso. Illustration at Kew (K001568512) with identification by M.A. Clements 6/9/1987 believed based on Colenso’s original material (i.e. K000827538) © RBG Kew
